# Genomic plasticity and rapid host switching promote the evolution of generalism in the zoonotic pathogen *Campylobacter*

**DOI:** 10.1101/080077

**Authors:** Dan J. Woodcock, Peter Krusche, Norval J. C. Strachan, Ken J. Forbes, Frederick M. Cohan, Guillaume Méric, Samuel K. Sheppard

## Abstract

Horizontal gene transfer accelerates bacterial adaptation to novel environments, allowing selection to act on genes that have evolved in multiple genetic backgrounds. This can lead to ecological specialization. However, little is known about how zoonotic bacteria maintain the ability to colonize multiple hosts whilst competing with specialists in the same niche. Here we develop a stochastic evolutionary model and show how genetic transfer of niche specifying genes and the opportunity for host transition can interact to promote the emergence of host generalist lineages of the zoonotic bacterium *Campylobacter*. Using a modelling approach we show that increasing levels of recombination enhance the efficiency with which selection can fix combinations of beneficial alleles, speeding adaptation. We then show how these predictions change in a multi-host system, with low levels of recombination, consistent with real *r/m* estimates, increasing the standing variation in the population, allowing a more effective response to changes in the selective landscape. Our analysis explains how observed gradients of host specialism and generalism can evolve in a multihost system through the transfer of ecologically important loci among coexisting strains.

## Introduction

Adaptation is typically thought to lead to gradual ecological specialization, in which populations progress towards an optimal phenotype. This can occur among competing organisms in sympatry, particularly when resources diversify or if the cost of maintaining homeostasis in different environmental conditions is high (Van Tienderen 1991). However, resource or host generalism are also widely observed in nature (Fried et al 2010, Kassen 2002, Woolhouse et al 2001) and it is generally accepted that natural environmental heterogeneity can promote the maintenance of phenotypic variation where it confers an ecological advantage (Kassen 2002, Van Tienderen 1991).

Host generalism is exhibited by some bacteria that infect multiple hosts resulting in important implications for the spread of disease from animals to humans. In some zoonotic bacteria, such as *Staphylococcus aureus,* livestock-associated lineages are largely host-restricted (Fitzgerald 2012), allowing the direction and time-scale of host transfer to be estimated by comparison of genotype information (Lowder et al 2009). In contrast, in *Escherichia coli* it is difficult to link ecological niche with genotype as isolates from all major phylogroups are represented in multiple isolate sources (Meric et al 2013). Other organisms including *Campylobacter jejuni* and *Salmonella enterica* (Baumler and Fang 2013) represent an intermediate between these. For example, comparison of *C. jejuni* isolates from various sources, by multilocus sequence typing (MLST) (Sheppard et al 2011) and whole genome sequencing (Dearlove et al 2015, Sheppard et al 2014), has shown evidence for host restricted lineages, found predominantly in one host species, as well as lineages commonly isolated from hosts as diverse as chickens, cattle and wild birds (Gripp et al 2011).

Out-competition by host specialists, might be expected to select against generalist *Campylobacter* lineages. However, generalists remain among the most common lineages in agricultural animals and are a major cause of human disease (Sheppard et al 2009). There are several factors that may be involved in the emergence of generalism as a successful strategy. First, the development, industrialisation and globalisation of livestock farming have created a vast open niche in which *Campylobacter* has expanded from pre-agriculture wild animal hosts. Second, factors associated with livestock husbandry and habitations promote close contact between different livestock species providing opportunities for *Campylobacter* transmission from one host to another. Third, *Campylobacter* is a highly recombinogenic organism (Wilson et al 2009) and lineages regularly acquire genetic elements through horizontal gene transfer (HGT). This genomic plasticity has the potential to introduce DNA segments, or whole genes, into the recipient genome, potentially conferring novel function.

The coexistence of *Campylobacter* lineages with generalist and specialist ecology remains poorly understood and little is known about the factors that promote the emergence and maintenance of lineages with distinct ecology (ecotypes). Here, using information on natural genomic variation in 130 *C. jejuni* genomes from chicken and cattle (Sheppard et al 2013b), as well as a computational model of bacterial evolution, we find that the maintenance of genetic variance of putative niche-specifying genes in the population can promote the emergence of generalism as seen in nature. By quantifying how resource competition, rapid host switching, and horizontal gene transfer interact to affect the variance in the population, we provide a generalised framework for considering the emergence of generalist ecotypes.

## Material and methods

### Bacterial genomes

A total of 130 *C. jejuni* and *C. coli* isolates were used in this study, including 87 representative strains sampled from chicken and cattle (Table S1). The genomes were previously published and isolates were described as belonging to clonal complexes on the basis of sharing 4 or more identical alleles at seven multilocus sequence typing loci (Sheppard et al 2013a, Sheppard et al 2013b). In total for this study, 30 genomes from the ST-21 clonal complex, 28 from the ST-45 clonal complex, 7 from the ST-353 clonal complex, 6 from the ST-206 clonal complex, 6 from the ST-61 clonal complex, and 5 from the ST-48 clonal complex were used. ST-353 complex isolates are known to be chicken-associated and ST-61 complex isolates to be cattle-associated, whereas ST-21, ST-45, ST-206 and ST-48 are host generalists (Sheppard et al 2014). Genomes were archived on a web-based platform system based on BIGSdb, which also implements analytic and sequence exporting tools (Jolley and Maiden 2010). An additional 75 genomes representing 74 different STs from *C. jejuni* and *C. coli* were used. These genomes were sequenced as part of other studies (Food Standards Scotland i-CaMPS-3 Contract S14054, DEFRA project OZ0625), and from the PubMLST Database (Jolley and Maiden 2010).

### Model input data

Using a gene-by-gene approach (Maiden et al 2013, Sheppard et al 2012), loci in the 130 genomes were identified by BLAST comparison to the *C. jejuni* strain NCTC11168 reference genome (Genbank accession number: AL111168) with a >70% nucleotide sequence identity on ≥50% of sequence considered sufficient to call a locus match, as in other studies (Meric et al 2014, Meric et al 2015, Pascoe et al 2015). A whole-genome MLST (Maiden et al 2013, Sheppard et al 2012) matrix was produced summarizing the presence/absence and allelic diversity at each locus in each genome, based upon these BLAST parameters. From this matrix, 1080 core genes were found to be shared by all cattle and chicken isolates from our dataset. The proportion of each allele at each locus was calculated in both cattle and chicken and then subtracted to identify alleles that were common in one group, but rare in the other. These were then summed these values at each locus to get the discriminative capacity of each locus for each host species. Loci in which the alleles segregated by host were considered as a proxy for niche-specifying genes in the model. While this was done in preference to simulating data, no inference is made based on the function or potential adaptive advantage conferred. A total of 5 putative niche-specifying genes per host (chicken and cattle) and 5 MLST genes picked at random (15 genes in total) were used as the input genotype for the model (**Table S2**). The inclusion of MLST loci (*aspA*, *uncA*, *pgm*, *glnA*, *gltA*) provides a reference for comparison.

### The Genome Evolution by Recombination and Mutation (GERM) model

The GERM model is a simplified representation of a bacterial population and associated processes, which allows us to simulate bacterial evolution *in silico* by tracking individual bacteria of variable genotypes as they are exposed to various selective environmental pressures. Furthermore, by simulating with a stochastic sampling algorithm, we can also incorporate some degree of the randomness inherent in natural populations, and hence investigate the importance of stochastic effects by performing simulations with the same initial population. Similar models have been proposed previously (Levin and Cornejo 2009), and our approach extends and builds on these models in a number of ways. In the model in this study, each individual bacterial cell is represented as a 15-locus genotype as described above. In a population of *N* cells, each cell *C* is described entirely by the alleles *a* at locus *j* which compose the genome, denoted *a_j_*, *j* = 1,2,…,15. Each allele at each locus is represented by an integer ranging from 0 to ∞. Basing our algorithm on real data, where there are approximately 20 possible alleles at each locus (20^15^ ≈ 3.27 × 10^19^ different combinations) presents computational challenges if we model the population using proportions of genotypes as in previous models (Levin and Cornejo 2009). To account for this we store the entire population at any one time and perform operations at the individual level. Working with the population directly, instead of adjusting proportions of STs, allows the investigation of the population dynamics at different population sizes, which is particularly pertinent after selective sweeps when the number in the population itself will drop as the population adapts to the new environment. The model incorporates six basic processes: mutation, recombination, resource consumption, cell death, cell division and host migration. Each process occurs once per generation and is stochastic, therefore occurring with a probability defined for each cell. These can be interpreted as rates per generation.

### Cell division, mutation and recombination

Mutation and recombination occur at the level of the individual locus and cell death and cell division occur at the individual bacterial cell level. With cell division, an identical copy of the cell that divides is added to the population, this occurs with probability *b*. Mutation occurs with probability *m* and, unlike existing models (Levin and Cornejo 2009) any allele that mutates is deemed to offer no selective advantage (the fitness of that allele is 0). Similarly, a recombination event occurs at probability *r* and an allele that recombines is assigned the value of another allele randomly chosen from those at the same locus within the current population. In natural systems, these processes are typically considered rare with upper rate estimates for homologous recombination of 10^−6^ per gene per generation (Wiedenbeck and Cohan 2011). However, these rate estimates can be affected by a number of factors (Barrick and Lenski 2013, Vos and Didelot 2009) and so typically in studies of bacterial evolution, the preferred measure of the magnitude of recombination is the relative frequency of recombination compared to mutation, the *r / m* ratio (Falush et al 2001, Fearnhead et al 2005, Feil 2004, Fraser et al 2005, Milkman and Bridges 1990). To quantify the effects of varying levels of homologous recombination on niche adaptation in the GERM model, we used *r/m* ratio as the ratio of rates at which alleles are substituted as a result of recombination and mutation. Partly for reasons of computational tractability, the GERM model simulates a simplification of the size and complexity of a natural system, and imposes enhanced selection against maladapted sequences types. Because of this, *r* and *m* rate estimates from natural populations are adjusted so that sequence types that arise in the population have a similar opportunity on average to proliferate in both the model and the natural environment and are not removed from the population by chance alone. Consistent with existing estimates from multiple bacterial species including *Campylobacter* (Vos and Didelot 2009), we run simulations at *r / m* ratios ranging from 0 to 100 corresponding to a mutation rate of 0.01, with recombination rates of 0, 0.001, 0.01, 0.1 and 1 to facilitate comparisons with natural populations.

### Fitness

In this study, host-specific alleles at niche specifying genes are considered to confer a fitness advantage to the cell in one or other host. The fitness of allele *a*, in a given host *h* is defined as *f* ^{*h*}^(*a*) and this reflects the fitness conferred by that allele to its environment, with 1 corresponding to a perfectly adapted allele conferring maximal fitness, ranging to an allele that provides no benefit to the survival of the cell and has 0 fitness. We then follow Levin (Levin and Cornejo 2009) and calculate the fitness of an individual cell as the sum of the allelic fitness values assuming that each allele contributes equally. The fitness function is therefore: 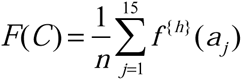. As a consequence, a different fitness landscape, defined by another host, affects the fitness of a cell through the fitnesses of its constituent alleles. It is by changing the fitness landscapes that host transition is simulated. Linkage disequilibrium is not factored into the model, although genes were selected from divergent genomic positions to limit linkage effects, and the model was not designed to test the specific function of individual genes. In nature, complex interactions between genes and the environment are likely to correspond to fitness function involving operations somewhere between additive and multiplicative (Phillips 2008) but less information on this is available for bacteria than in the more well studied diploid scenario.

### Resource

The fate of a bacterium in the GERM model is not only dependent on fitness, but also on the availability of resources. This introduces a dynamic relationship between the fate of cells, their fitness, and the population size, and confers a soft carrying capacity for a niche dependence on the interplay between these aspects. Resource is modeled as a generic entity for which no distinction is made for the type of resource in a given niche, and the affinity for the consumption of a resource by a bacterium is independent of genotype. For each cell, the chance of using a resource occurs with probability *u*^+^, multiplied by the amount of available resource. If a cell is already using a resource, it finishes with probability *u*^−^ in which case the resource is then consumed and is not returned to the environment. A resource is generated with constant probability *g* and is added to the pool of available resource. As such, for any given death in the population, there is a probability that it is caused either by a lack of fitness, or the inability to find and utilize resource. For a given fitness, the chance of death due to fitness or resources changes as the number of cells rises. For very small populations, the cause of death is predominantly fitness-related as resources are abundant. As the population increases, and the chance of finding a resource diminishes, resource-related death quickly starts to influence the fate of the cell but then plateaus as the population approaches carrying capacity. This nonlinear relationship is because, when unconstrained, the population will double in size during the course of a generation, where the resources will only increase by a fixed amount each time, regardless of population size. Conversely, as fitness increases, the proportion of deaths caused by fitness decreases to the point that resource related death becomes dominant. This is important as the mechanism of death is different in each case: in fitness-related death, the probability of death is inversely proportional to the fitness. However, for resource-related death, the chance of death is independent of fitness and occurs entirely at random. As such, when resources become scarce, the fitness of the members of a population becomes largely less important to their fate. Therefore, resource availability is fundamental to the population dynamics, as well as suppressing the speed of convergence to the optimal genotype and maintaining genetic variance.

### Cell death

Cell death occurs in two stages, fitness-related death and resource-related death. Fitness-related death is dependent on a fitness function, as discussed above and detailed further the supplementary material, which provides a probability of dying per generation for a given genotype. Resource-related death can happen only when a cell is not using a resource and occurs with probability *d* regardless of fitness. In both cases, the cell is removed from the population. The algorithm also incorporates a host transition or a selective sweep by switching between fitness functions. In this case, the resources can also be reset (although those already using a resource will continue as before). Further details of the model and stochastic simulation are included in the supplementary material.

### Simulation

To account for the randomness inherent in the process of evolution, we simulated the system with a variant of a stochastic sampling algorithm (SSA) (Gillespie 1977), involving Monte Carlo decision processes, was used for simulating the system. The number of cells required to represent a natural population is extremely large, therefore some constraints were imposed to ensure computational tractability. We sampled from Poisson distributions and binomial distributions when the quantities in question were unbounded and bounded respectively. For simplicity, we set the probability of a cell division event *b* to be equal to 1 so that every iteration can be interpreted as a generation. For mutation and recombination we imposed a constraint that each of these events can occur only once every time step. There were ten time steps per generation. The algorithm iterated through the five processes in order of resource allocation, mutation, recombination, cell death and cell division. We use three fitness functions, one each from chicken and cow, which we subsequently refer to as “Host 1” and “Host 2” respectively, and one derived using data from both hosts referred to as the composite fitness function. The composite fitness function is used to initialize the simulation and allow a burn-in period where the genetic composition to arrange themselves in a way that favors neither host. The algorithms start in using the composite fitness function to allow the algorithm to burn-in and deplete unfit allele combinations without purging alleles from one of the hosts. When the population has recovered and reached equilibrium with resource generation, we switch the fitness function to one of the single hosts. This occurred after 200 generations. We subsequently alternated the fitness function between chicken and cattle with the frequency defined by the user. This corresponds to the movement of the entire population to the new host, a simplification of the more realistic process in which bacteria would move to one of several distinct and concurrently evolving ecosystems. This layer of abstraction was partially chosen for computational tractability, but also to aid in the interpretation of results as the number of stochastic events would increase exponentially as the model complexity increases which would potentially dominate any patterns in our results. As such, this constraint is suitable for the scope of this study, particularly as our model is predicated on *Campylobacter* populations which are often transmitted *en masse* through stool. To initialize the algorithm, we generated a population of 50 million cells consisting of alleles found in the data set, randomly assigned throughout the population in a uniform manner rather than weighted by their abundance in the data set. The same initial population was used for every simulation run. This way, the populations and processes can be kept identical, and so that the only differences between runs with the same parameters were stochastic effects.

Two experiments were carried out at a range of recombination to mutation ratios (0-100).

1. Long term adaptation followed by host switch – after the first composite burn in, a switch to host 2 was performed. This was run for 1000 generations to simulate long term colonization and adaptation and was then followed by a switch to host 1.
2. Rapid host switching – after the composite burn in – rapid host switching (every 200 generations) was performed.

### Host generalism and phylogeny

We used 75 genomes from isolates representing 74 *C. jejuni* and *C. coli* STs, from 30239 pubMLST isolate records (http://pubmlst.org/campylobacter/), to investigate the degree of host generalism among different lineages. From these, concatenated gene-by-gene alignments of 595 core genes (Sheppard et al 2013b), constructed as previously published (Meric et al 2014, Meric et al 2015), were used to infer phylogenetic relationships. A tree was constructed using the neighbor-joining algorithm and each ST was labelled with the number of distinct hosts from which it has been isolated, based on data submitted to the pubMLST database. The 74 STs were reported to have been isolated from 20 distinct animal hosts, including humans. The maximum number of hosts associated with a single ST was 12 (ST-45) and the minimum was 1 (ST-58). A tree and the heatmap representing the number of hosts, was prepared and visualized in Evolview (Zhang et al 2012). The mean host species richness score was correlated with the genetic distance (derived from the number of SNPs) from the tip of the tree to the first branching point for each of the isolates.

## Results

### Long term adaptation promotes specialism but recombination enhances colonization in a subsequent host transition

The effect of homologous recombination was characterized in 200 independent population GERM simulations, with a transition from the composite niche to Host 1 after 200 generations, followed by a transition to Host 2 after another 1000 generations. Simulations were performed at five *r /m* ratios: 0; 0.1; 1; 10; 100. In all simulations, the mean number of cells decreased sharply from the initial condition and then recovered to approach an equilibrium level between birth and death just before the transition to Host 1 (Figure 1). In the composite niche, level of proliferation was proportional to the rate of recombination with concomitant increase in mean fitness and population genetic variance (Figures 1C and 1D). This is because higher recombination rates result in greater genetic variance, and so by Fisher’s fundamental theorem of natural selection (Fischer 1930), the rate of increase in the mean fitness will be greater and the population will thrive. Following the first host transition (Figure 1A, numeral I), there was a brief increase in the population due to increased resource availability, but in all cases the mean number of cells quickly returned to the equilibrium state where it continued until the next host transition (numeral II), with the mean number of cells ordered largely as before, with the number of cells increasing with recombination rate (Figure 1C). After the transition to Host 2, the populations were decimated in all cases, as the alleles which would have conveyed enhanced fitness in Host 2 have been purged from the population, meaning that the bacteria are ill-equipped to survive in the new host and so they will die. Almost all of the populations died out in the low recombining groups (*r*/*m*=0 and 0.1), a large proportion died out in the intermediate group (*r*/*m*=1), with the greatest number of surviving populations in the highly recombining groups.

**Figure 1.**
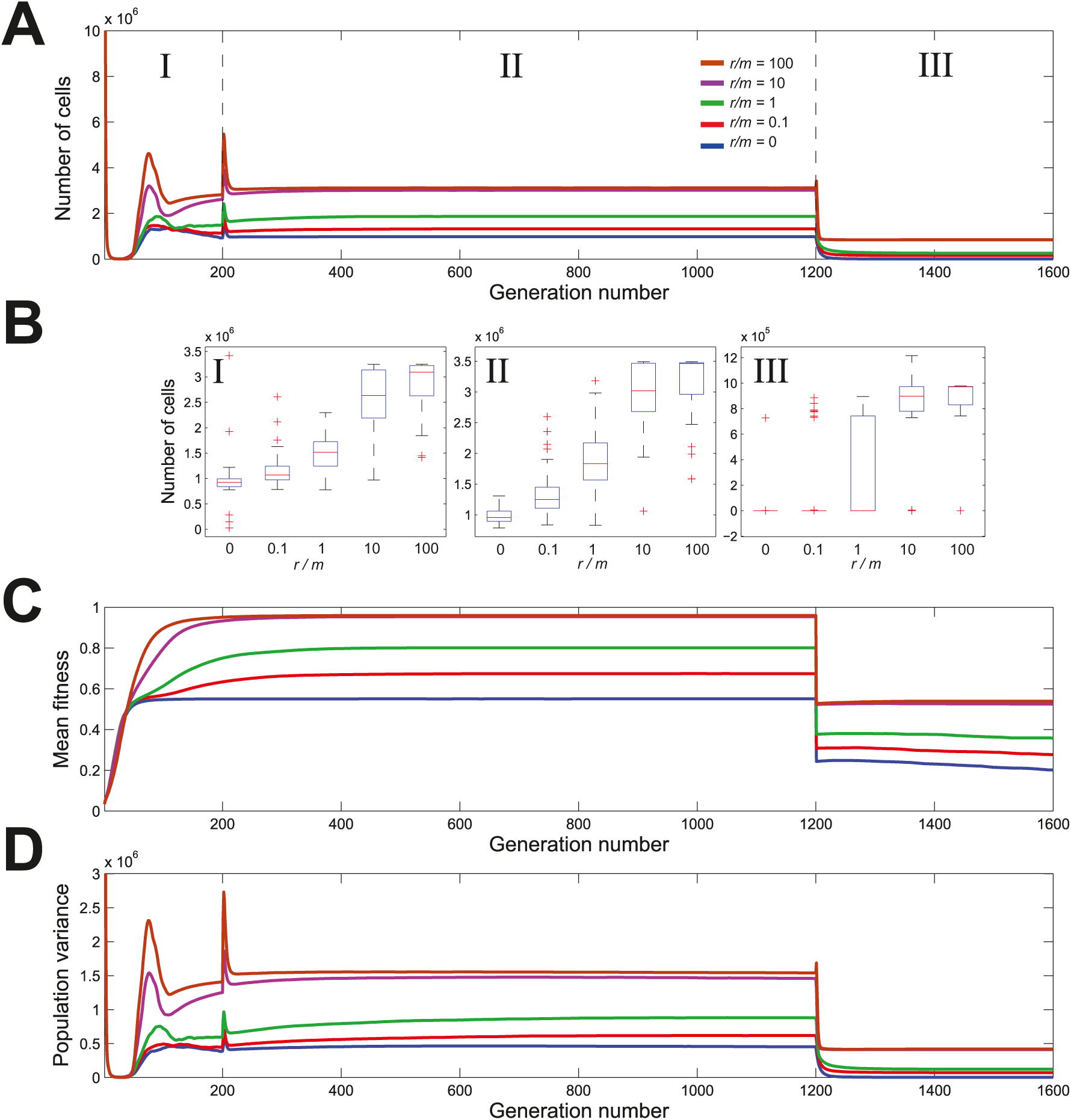
Long term host adaptation followed by a host switch with different recombination rates. The number of cells (A), population mean fitness (C) and population variance (D) were monitored in 200 independent simulations performed at recombination to mutation ratios of: *r/m*=0 (blue), *r/m*=0.1 (red), *r/m*=1 (green), *r/m*=10 (magenta) and *r/m*=100 (brown). Model parameters were different in phases I, II and III. Phase I corresponds to a composite niche no fitness-related selection. The transition from I to II corresponds to the addition of selective pressure favoring genes specifying adaptation to host 1, and the transition from phase II to III, corresponds to a single host transition, with a change to selective pressures favoring genes specifying adaptation to host 2. Panel B shows the number of cells at the end of every phase for populations with different recombination rates.

### Intermediate recombination rates enhance population mean fitness after multiple rapid host transitions

As in the single host transition model, the mean number of cells in the composite niche reached equilibrium levels ordered by their recombination rate (Figure 2A) and consistent with their mean fitness levels (Figure 2B). At the end of growth in the composite niche, the intermediate and high recombination rates (*r/m*=1, 10 and 100) have a similar ability to survive, with the highest recombination level displaying high variance (**Figure 2C**). After the composite niche a number of host transitions were simulated where the mean number of cells shifted between two equilibrium states depending on the host species. The mean number of cells was always higher in Host 1 because some alleles conferring increased fitness in Host 2 will inevitably be purged from the populations in the Host 1 niche. Reversing the species order had the same effect for Host 2. At the end of the last Host 2 growth cycle all the recombining populations (*r*/*m*>0) had a similar mean number of cells, but the variance differed, with the smallest variance at *r/m*=1 and *r/m*=10. Similarly, in the final Host 1 niche, we can see in Figure 2A that the intermediate recombination rate (*r/m*=1) is associated with the highest mean number of cells. This shows that in contrast to the single host transition model, recombination at a high level is not advantageous under repeated host switching.

**Figure 2.**
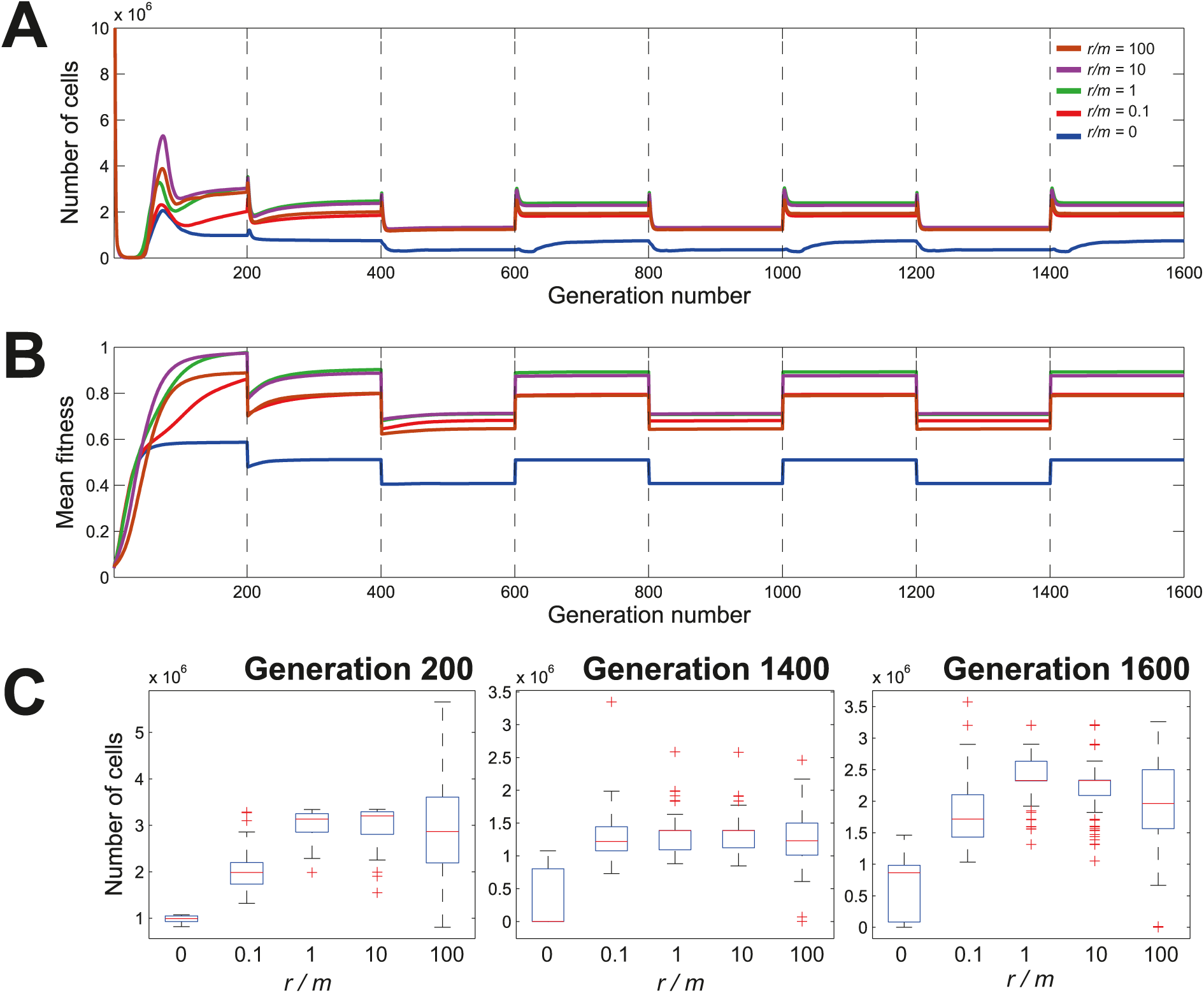
Rapid multiple host transitions with different recombination rates. Number of cells (A) and mean fitness (B) for recombination to mutation ratios of *r/m*=0 (blue), *r/m* =0.1 (red), *r/m* =1 (green), *r/m* =10 (magenta) and *r/m* =100 (brown) as they progress through several niche transfers (broken lines), for which selective pressures are alternatively imposed to favor genes specifying adaptation to one or the other host. (C) Population growth distribution at various recombination rates at generations 200 (transition from composite niche to the first host), 1400 (last transition growth phase in host 2) and 1600 (end of simulation after 6 host switches).

### Emergence of host generalist strategy as a consequence of frequent host switches

We used a Dirichlet process clustering algorithm (Kurihara et al 2007) on all simulations to identify characteristic profiles of population dynamics for the different recombination rates (Figure 3). Three broad population dynamic profiles were observed: (i) populations that were primarily adapted to Host 1 (Clusters 1-5); (ii) populations that were primarily adapted to Host 2 (Clusters 7-9); (iii) populations that were adapted to both niches (Cluster 6). This is consistent with a classification as a specialist for either species, or as a generalist. The membership of these 3 adaptation profile types relates to recombination rate (Table 1). Simulations with no recombination were predominantly found in Cluster 2, with a substantial amount found in other clusters. This is to be expected as the outcome was driven entirely by stochasticity acting on the population and so genes were purged almost at random in the composite niche, yielding a set of outcomes which were maintained during the host transitions as more alleles were lost. In contrast, it can be seen that simulations of all of the recombining populations are predominantly found in Cluster 4, which is a Host 1 specialist cluster, albeit with a relatively high equilibrium population number during the Host 2 niche compared to the other Host 1 specialist clusters. In Cluster 4, it can be seen that an *r/m*=1 gives the greatest occupancy, at 91.8%, explaining the high numbers of cells seen in Figure 2. The membership of the generalist cluster, Cluster 6, is also represented across all recombination rates, with the highest percentage coming from a relatively low recombination rate (*r/m*=0.1).

**Figure 3.**
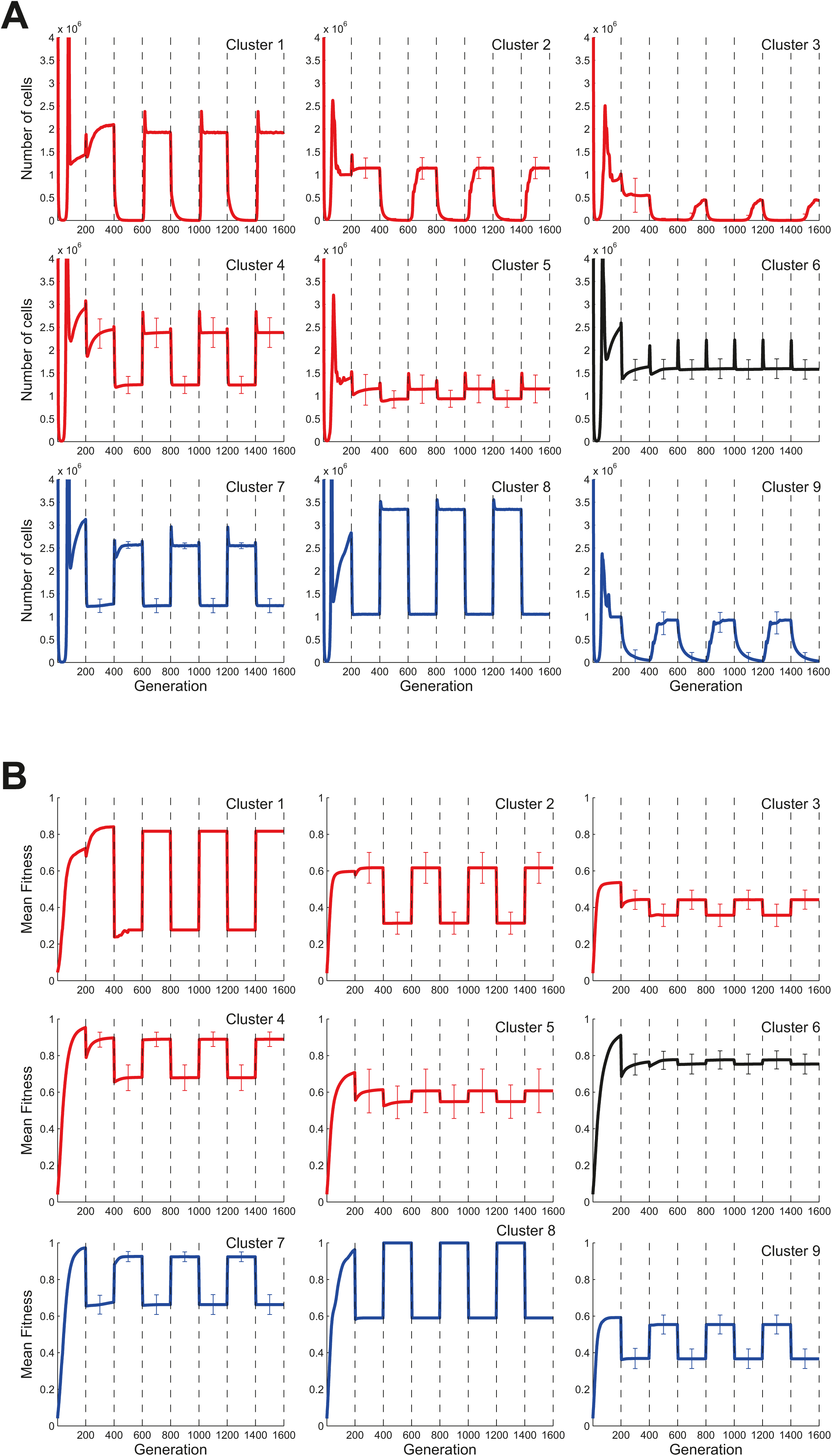
Population dynamic profile clusters. Population profiles for all simulates at tested recombination rates. Nine distinct profile clusters were identified and the mean number of cells (A) and the mean fitness (B) is shown for example simulations for profile cluster. Black broken lines indicate a host transition where selective pressures switch in favour of the other host. Error bars are given at the midpoint of each niche and correspond to one standard deviation. Ecological groups were inferred from the profiles, with specialist groups for Host 1 (red line) and Host 2 (blue line) as well as a generalist group (black line).

**Table 1:**
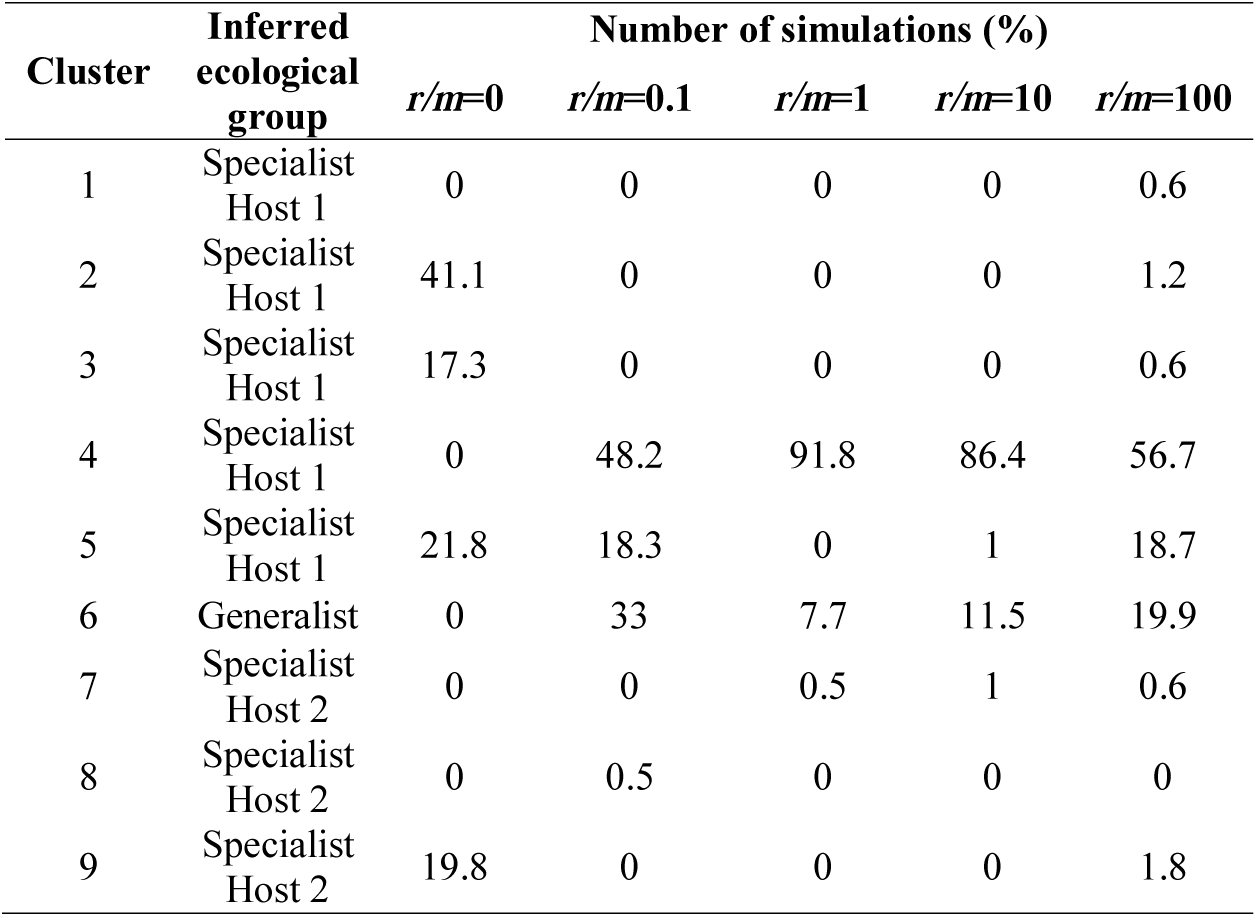
**Proportion of the representative clusters of population profiles at various recombination to mutation ratios**. For each ratio (*r / m*), the percentage of simulations that followed particular representative patterns (clusters 1-9 from Figure 3) from a total of 200 simulations were indicated. Ecological groups were inferred from Figure 3.

### The gradient of host generalism is mirrored in natural *Campylobacter* populations

The degree of host specialism and generalism in *in silco*, resulting from model simulations, was compared to data from natural *Campylobacter* populations. Mapping isolation source data of 30239 *Campylobacter* STs within the PubMLST database (Jolley and Maiden 2010) to a core genome neighbor joining tree of the 74 common *C. jejuni* and *C. coli* STs revealed that few STs demonstrate absolute generalism or specialism (Figure 4a). Rather there is a gradient ranging from STs predominantly found in one host to those frequently isolated from multiple host sources (Figure 4b). The clustering of isolate pairs, with shared host source richness, was estimated by correlating nucleotide divergence in the core genome with the number of hosts (Figure 4c) and by a randomisation/ permutation test which showed p<0.000001 (data not shown). STs on the phylogeny were located close to STs with similar host species richness suggesting that there is some evolutionary signal which determines the likelihood of an ST’s degree of specialism.

**Figure 4.**
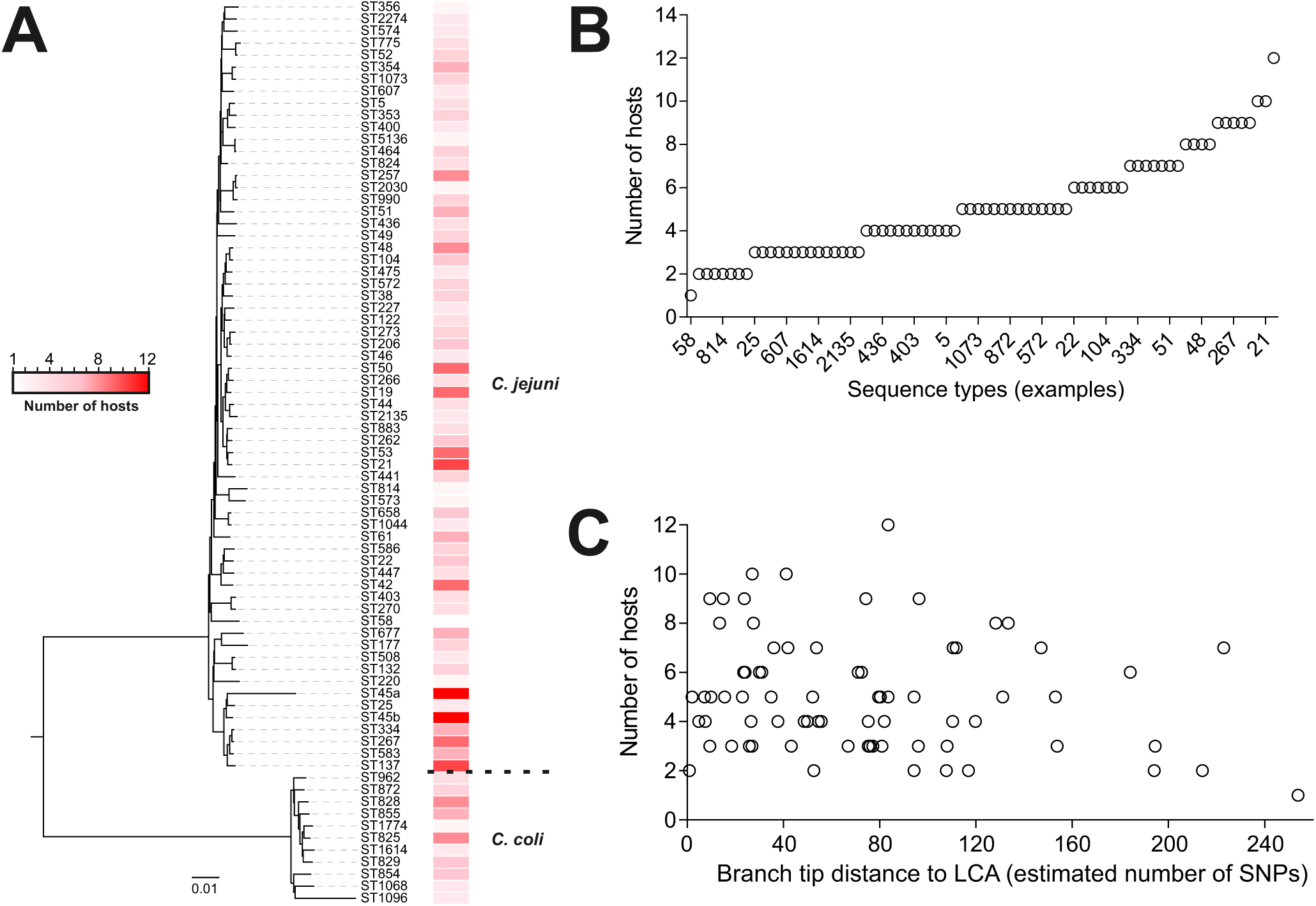
Phylogeny of generalist and specialist *Campylobacter* lineages. (A) Phylogenetic tree and isolation source of 74 common C. jejuni and C. coli sequence types (STs). The tree was constructed from a concatenated gene-by-gene alignment of 595 core genes, using the neighbor joining (NJ) algorithm. The heatmap represents the number of hosts in which particular STs were isolated, based on analysis of 30239 pubMLST isolate records. The scale bar represents the number of substitutions per site; (B) Quantification of the variation in number of hosts for all lineages shown in the tree, highlighting a gradient between host specialism and generalism; (C) Comparison of clustering on the tree, calculated as the estimated number of SNP corresponding to the average branch tip distance to the last common ancestor (LCA) with the number of hosts of isolation of each examined ST. There is no correlation between the two datasets (linear regression; r²=0.037).

## Discussion

The GERM model provides a context for considering how genome plasticity may influence the proliferation of *Campylobacter* in a multihost environment. In simple simulations, rapid acquisition of niche-specifying genes promoted better colonization in a new host. This is consistent with the Fisher–Muller evolutionary model (Fisher 1930, Muller 1932) where recombination functions to bring together fit alleles, which would otherwise compete for fixation in the population, into a single lineage speeding the overall increase in population mean fitness (Gerrish and Lenski 1998). In line with classical population genetic theory for sex (Barton and Charlesworth 1998, Felsenstein 1974, Weismann 1904), the efficiency of selection on bacteria is enhanced by this shuffling of alleles. Therefore, the population with the highest recombination rate will expand to fill the niche more rapidly after a genetic bottleneck. This demonstrates a clear short-term adaptive benefit to rapid recombination, but does not explain why most bacteria recombine at low rates in nature.

Where survival is predominantly influenced by a few genetic determinants, for example the acquisition of essential antibiotic resistance genes (Spratt et al 1989), high recombination rates would be favoured. However, this is an unusually simple evolutionary scenario and bacterial habitats comprise numerous interacting selective pressures. Because increased genetic variation leads to faster adaptation (Fisher 1930), the potential for the population to survive future genetic bottlenecks is related to the fitness variance. In populations that recombine at a low rate, a relatively high fitness variance is often maintained. Therefore, if the species is likely to encounter frequent environmental changes, such as host switches, it may be beneficial to have a lower recombination rate than would be optimal in the Fisher-Muller model.

In nature, each niche will have an associated carrying capacity. As a species expands to approach this carrying capacity, competition will inevitably increase, meaning that the influence of external factors will be of increased importance to the fate of the organism. To avoid this competition, an organism could adapt to a new less competitive niche, consistent with the ‘tangled bank hypothesis’ (Bell 1982, Doebeli 1996, Koella 1988) for the evolution of sexual reproduction. In our competitive model, fitness variance is higher in populations with low recombination rates, providing a competitive advantage under the tangled bank hypothesis that would facilitate a transition to a new niche. This provides an explanation for the low recombination rates observed in some natural bacterial populations, and explains the presence of multiple lineages as infrequent recombination will allow the uptake of adaptive genes but may be too infrequent to prevent adaptive divergence between lineages (Wiedenbeck and Cohan 2011).

Based on model simulations, the most favorable recombination to mutation ratio for promoting *Campylobacter* survival in the new niche whilst maintaining fitness variance within the population was *r* / *m* =0.1-1 which is comparable to that calculated in natural *Campylobacter* populations (*r* / *m* =0.44) (13). In multiple niche transition simulations, at intermediate recombination rates (*r / m =* 1), many populations did not completely die out, but resisted the introduction to a novel host recovering after a few passages, as seen by population size and mean fitness increases over time. This provides a context for considering the balance between rapid adaptation, mediated by recombination, and maintenance of genetic variance, allowing each population to survive in both host environments over time (Figure 1 and Figure 2). Absolute host specialists, which went extinct in the second host, were uncommon, and most populations demonstrated some capacity to survive in both hosts.

In most cases, simulated genotypes were more successful in one or other niche. This is consistent with evidence from natural populations where lineages such as the ST-257 and ST-61 clonal complexes are predominantly associated with chicken and cattle respectively but are also isolated from both niches (Sheppard et al 2014). However, in some multihost simulations, populations emerged that were affected very little by the host switches (Figure 3). These populations can be considered true ecological generalists, comparable to the ST-21 and ST-45 clonal complexes that are regularly isolated from chickens, cattle and other hosts (Sheppard et al 2014).

Ecological specialism and generalism have been well described in animals. For example, the Giant Panda, *Ailuropoda melanoleuca*, is a paradigm of specialism, confined to six isolated mountain ranges in south-central China, where bamboo comprises 99% of its diet (Lü et al 2008), while the American Black Bear, *Ursus americanus*, is a generalist, opportunist omnivore with a broad range including temperate and boreal forests in northern Canada and subtropical areas of Mexico (Garshelis et al 2008). Specialization is a potentially precarious strategy as change to the environment can cause extinction if organisms are unable to move between niches or hosts. Consistent with this, generalist lineages would be expected to be older, preceding specialists which cluster closer to the tips of the phylogenetic tree (Stireman 2005). In *Campylobacter,* there was no correlation between tree tip length and number of hosts suggesting similar evolutionary timescales for specialist and generalist STs. This contrast with studies of metazoans may, in part, be explained by the ability of *Campylobacter* to rapidly acquire niche-specifying elements leading to rapid adaptation of multiple lineages.

In addition to genomic plasticity, the scale of environmental variation can act on the type of ecological strategy observed among the inhabitants (Futuyma and Moreno 1988, Kassen 2002). For example, in a highly stratified environment, adaptation may occur early with traits becoming fixed, whereas in a graduated environment individuals may be more likely to show a reversible phenotype response (Kassen 2002). Specific niches of different bacterial lineages may be less well defined than for some animal species, but isolation source data from the PubMLST database indicates that *Campylobacter* STs show a gradient of host generalism. This is influenced by the opportunity to colonize the new niche and the capacity to survive the transition. Consistent with the findings in this study, rapid host transitions provide an opportunity for the proliferation of organisms with varying levels of host specialization including generalists that are capable of proliferating in more than one host.

In this study, by combining data from natural bacterial populations and a selection driven computer modeling, we simulated and predicted the evolution of host association strategies in the zoonotic pathogen *Campylobacter*. In practice, bacteria in natural populations usually exist in complex changeable ecosystems with numerous selection pressures. Here we show that recombination allows a more rapid response after a genetic bottleneck, as in a host transition, by increasing the efficiency with which selection can fix combinations of beneficial alleles. Furthermore, in a dynamic setting of host switching, recombination rate was observed to be a key factor in the colonization and maintenance in multiple niches. Livestock in modern intensive agricultural systems are different to ancestral host populations in numerous ways associated with diet and stocking density. The implications of this are potentially significant as, under conditions favoring rapid host switching, the emergence of host generalist zoonotic pathogens can be simulated. Our model therefore provides a context for considering how recombining bacteria, such as *Campylobacter,* could evolve to meet the challenges of anthropogenic environmental change. This could promote the emergence of multi-host pathogens and increase their capacity to overcome deliberate human interventions.

## Acknowledgements

This work was supported by the Biotechnology and Biological Sciences Research Council (BBSRC) grant BB/I02464X/1, the Medical Research Council (MRC) grants MR/M501608/1 and MR/L015080/1, and the Wellcome Trust grant 088786/C/09/Z. GM was supported by a NISCHR Health Research Fellowship (HF-14-13).

